# Discovery of genes encoding a Streptolysin S-like toxin biosynthetic cluster in a select highly pathogenic methicillin resistant *Staphylococcus aureus* JKD6159 strain

**DOI:** 10.1101/752204

**Authors:** Trevor Kane, Katelyn E. Carothers, Yunjuan Bao, Won-Sik Yeo, Taeok Bae, Claudia Park, Francisco R. Fields, Henry M. Vu, Daniel E. Hammers, Jessica N. Ross, Victoria A. Ploplis, Francis J. Castellino, Shaun W. Lee

## Abstract

**Background:** *Staphylococcus aureus* (*S. aureus*) is a major human pathogen owing to its arsenal of virulence factors, as well as its acquisition of multi-antibiotic resistance. Here we report the identification of a Streptolysin S (SLS) like biosynthetic gene cluster in a highly virulent community-acquired methicillin resistant *S. aureus* (MRSA) isolate, JKD6159. Examination of the SLS-like gene cluster in JKD6159 shows significant homology and gene organization to the SLS-associated biosynthetic gene (sag) cluster responsible for the production of the major hemolysin SLS in Group A *Streptococcus*.

**Results:** We took a comprehensive approach to elucidating the putative role of the sag gene cluster in JKD6159 by constructing a mutant in which one of the biosynthesis genes (sagB homologue) was deleted in the parent JKD6159 strain. Assays to evaluate bacterial gene regulation, biofilm formation, antimicrobial activity, as well as complete host cell response profile and comparative *in vivo* infections in Balb/Cj mice were conducted.

**Conclusions:** Although no significant phenotypic changes were observed in our assays, we postulate that the SLS-like toxin produced by this strain of *S. aureus* may be a highly specialized virulence factor utilized in specific environments for selective advantage; studies to better understand the role of this newly discovered virulence factor in *S. aureus* warrant further investigation.

## Background

*Staphylococcus aureus* (*S. aureus*) is a pathogenic bacterium that is capable of causing severe human diseases including skin infections, pneumonia, and blood sepsis [1–4]. Of particular concern is the ability of *S. aureus* to acquire resistance to antibiotics. *S. aureus* presents as resistant to the frontline β-lactam antibiotic methicillin in roughly half of all infections in the United States [5]. Based on where the infection is acquired, methicillin-resistant *S. aureus* (MRSA) is classified as community acquired MRSA (CA-MRSA) or hospital acquired MRSA (HA-MRSA), accounting for some 84% and 16%, respectively, of clinical isolates in the United States [4]. CA-MRSA tends to be highly virulent and can produce toxins that are exclusive to CA-MRSA such as Panton Valentine leucocidin [6]. *S. aureus* typically encodes a wide range of virulence factors including protein A, enterotoxins, and α-toxin among others. This work examines *S. aureus* JKD6159, a highly virulent strain of CA-MRSA from Australia. The JKD6159 strain has become the dominant strain of MRSA found in Australia despite the initial isolation of the strain only 15 years ago [7, 8]. Our initial bioinformatic survey of this CA-MRSA strain indicated that this strain possesses a biosynthetic gene cluster homologous to the Streptolysin S associated gene (*sag*) cluster found in a related human pathogen, Group A *Streptococcus* (GAS). The *sag* cluster in GAS encodes the biosynthetic machinery required for production of Streptolysin S (SLS), a potent hemolysin and a major virulence factor in GAS pathogenesis [9, 10].

The Streptolysin S-associated gene (*sag*) cluster was initially identified by Nizet and colleagues using transposon mutagenesis in GAS [9, 10]. The operon responsible for SLS production was found to consist of nine genes that when disrupted individually resulted in the loss of the hemolytic phenotype [9]. SLS is the most well-studied member of a family of post-translationally modified peptides known as linear azole containing peptides (LAPs) [11–13]. These peptides are ribosomally synthesized and subsequently post-translationally modified *via* the installation of azole heterocycles on cysteine, serine, or threonine residues [12]. Peptides in the LAP family demonstrate a range of functions, including toxins such as SLS, and antimicrobial peptides such as Microcin B17 [9, 10, 14].

Although SLS has long been studied as a major lysin of red blood cells, recent studies have demonstrated that SLS may have specific and multiple cellular targets [15–20]. SLS has been identified as major contributing factor in successful translocation of Group A *Streptococcus* across the epithelial barrier during the early stages of infection [17]. Specifically, SLS facilitated breakdown of intracellular junctions through recruitment of the cysteine protease calpain, which cleaves host proteins such as occludin and E-cadherin [17]. SLS has also been implicated in the induction of programmed cell death in macrophages and neutrophils as well as inhibition of neutrophil recruitment during infection [18–20]. SLS has also been shown to induce several specific host cell signaling changes in human keratinocytes. Infection of HaCaT cells with wild-type and SLS-deficient GAS results in differential phosphorylation of p38 MAPK and NF-κB [15]. These recent insights into additional roles for pore-forming hemolysins highlight the potential contribution of SLS and SLS-like toxins in bacterial pathogenesis at the cellular level under conditions that more closely mimic the infection process.

Following our initial discovery of a sag-like biosynthetic gene cluster in a highly virulent methicillin resistant strain of *S. aureus* JKD6159, we assessed a possible role of the *sag-like* gene cluster in this *S. aureus* isolate, by using a comprehensive series of tests to evaluate a potential functional role for this genetic cluster using the wt and a genetic mutant of JKD6159 where the SLS-like biosynthetic gene *sagB* was inactivated. The wt JKD6159 strain and the *sagB* isogenic mutant displayed no significant differences in hemolysis, antibacterial activity, biofilm formation, host cytotoxicity or *in vivo* pathology. However, we noted changes in host cell signaling profiles between wt and the *sagB* mutant using a cell culture infection model, as well as differences in proteomic profiles between wt JKD6159 strain and the *sagB* isogenic mutants. We propose that further studies are needed to better define context-dependent conditions under which the SLS-like toxin will contribute to *S. aureus* pathogenesis and fitness in this highly virulent patient isolate.

## Results

### Identification of the *sag* cluster in *S. aureus* JKD6159

To identify the presence of a sag-like biosynthetic gene cluster in possible methicillin resistant strain human isolates of *S. aureus*, the gene and protein sequence from GAS strain M1 *sagB* were used to query non-redundant gene or protein sequences in NCBI using BLAST. We observed a match on the *S. aureus* strain JKD6159 containing a *sag* cluster with architecture similar to the *sag* cluster in GAS (Figure 1A). The gene cluster observed in the *S. aureus* strain contained all of the components of the *sag* cluster as found in GAS including the *sagB, sagC*, and *sagD* modifying enzymes as well as ABC transporters (Figure 1A). We also observed several putative small open reading frames in the vicinity of the modifying genes that could serve as the protoxin gene (Figure 1A). Analysis of the GC content in the gene cluster indicates it was likely horizontally acquired and integrated between the *gnd* gene and the *gloxalase* gene (Figure 1B). To assess the role of the *sag-like* gene cluster in *S. aureus* JKD6159, we generated a *sagB* isogenic mutant in JKD6159, which displayed no growth defects compared to the wildtype strain (Additional Files: Figure S1). We performed a comprehensive series of tests to evaluate a potential functional role for this genetic cluster.

**Figure 1:**
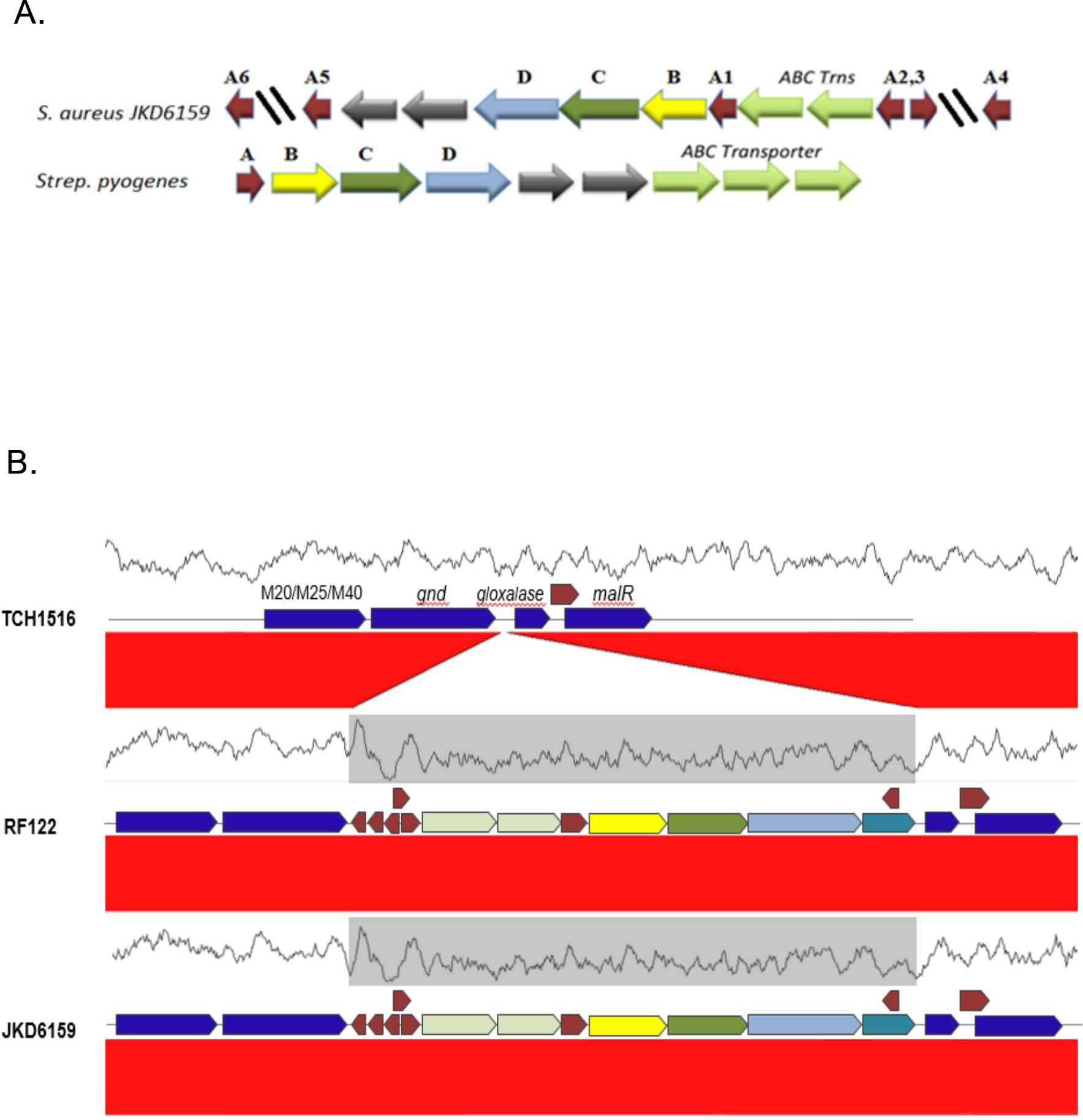
Presence of the *sag*-like gene cluster in *S. aureus* strain JKD6159. (A). Genetic organization of the *sag*-like gene cluster in strain JKD6159. A1-A6 denote putative protoxin open reading frames. The Streptolysin S biosynthetic cluster is shown below for comparison. Homologous genes are coded by colors, *sagB*-yellow, *sagC*green, *sagD*-blue. Potential protoxin genes are indicated in red. (B). GC analysis of the *sag* like cluster in JKD6159 and bovine isolate RF122. USA300 TCH1516 was used as the outgroup as it lacks the *sag* cluster. GC profile similarity between the RF122 strain and the JKD6159 strain are denoted in gray.

### Sheep blood hemolysis assays

As hemolysis is one of the hallmarks of SLS [9, 10], the JKD6159 *sag* cluster was hypothesized to confer production of a toxin with hemolytic activity. Prior work examining the SLS toxin in GAS showed that disruption of any of the nine genes in the *sag* cluster results in loss of hemolytic ability [9]. An isogenic deletion in the *sagB* gene was thus generated in JKD6159. Defibrinated sheep blood was used to examine the hemolytic potential of the conditioned media from the wt and Δ*sagB* JKD6159 strain. 25% conditioned media from overnight growth was sterilized and added to sheep blood and incubated at 37°C for the indicated time (Figure 2). Hemolysis levels were normalized to a TSB negative control and a 0.1% Triton X-100 positive control. No significant difference in hemolysis was observed between the wt and Δ*sagB* JKD 6159 strain (Figure 2).

**Figure 2:**
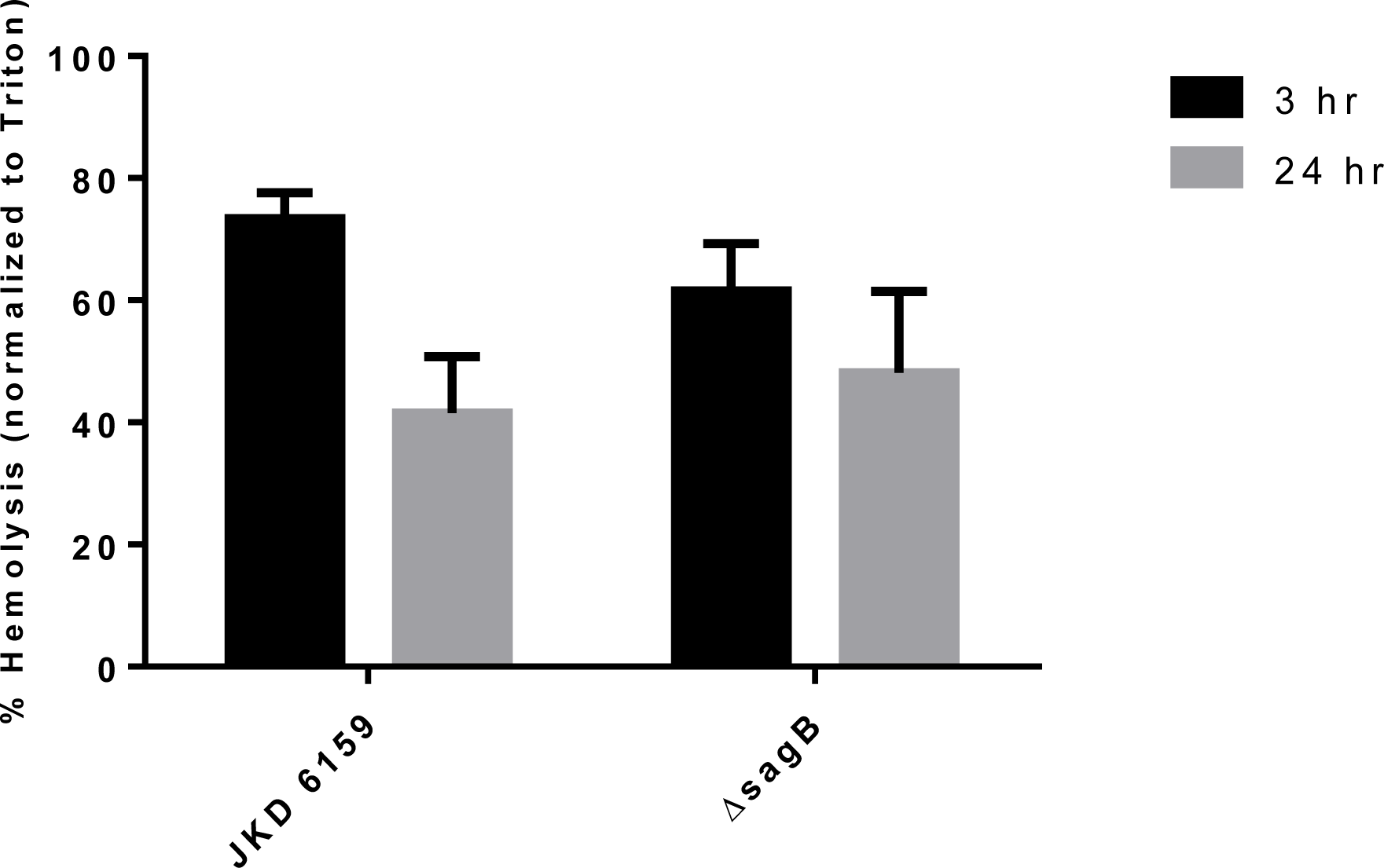
Red blood cell hemolytic activity of supernatants from *S. aureus* strain JKD6159 and isogenic Δ*sagB* mutant. Hemolysis was normalized to Triton X-100 activity controls. RBCs are incubated with 25% v/v conditioned media from wt or Δ*sagB* strains of *S. aureus* for times indicated.

### Cytotoxicity assays

Streptolysin S has been shown to promote cytotoxicity in various cell lines, including keratinocytes [15]. Cytotoxicity of the wt and Δ*sagB* JKD6159 strains were assessed by model infection of a keratinocyte cell line (HaCaT) at an MOI of 50 using a transwell infection system. Transwells were utilized to minimize contact-dependent effects that could potentially contribute to host cell cytotoxicity readouts. As the only difference in soluble factors between the wt and Δ*sagB* JKD6159 is the putative *sag-*encoded toxin, we can compare the readouts of cytotoxicity and attribute to the putative toxin produced by the *sag* cluster in JKD6159. No significant differences in cytotoxicity were observed (Figure 3).

**Figure 3:**
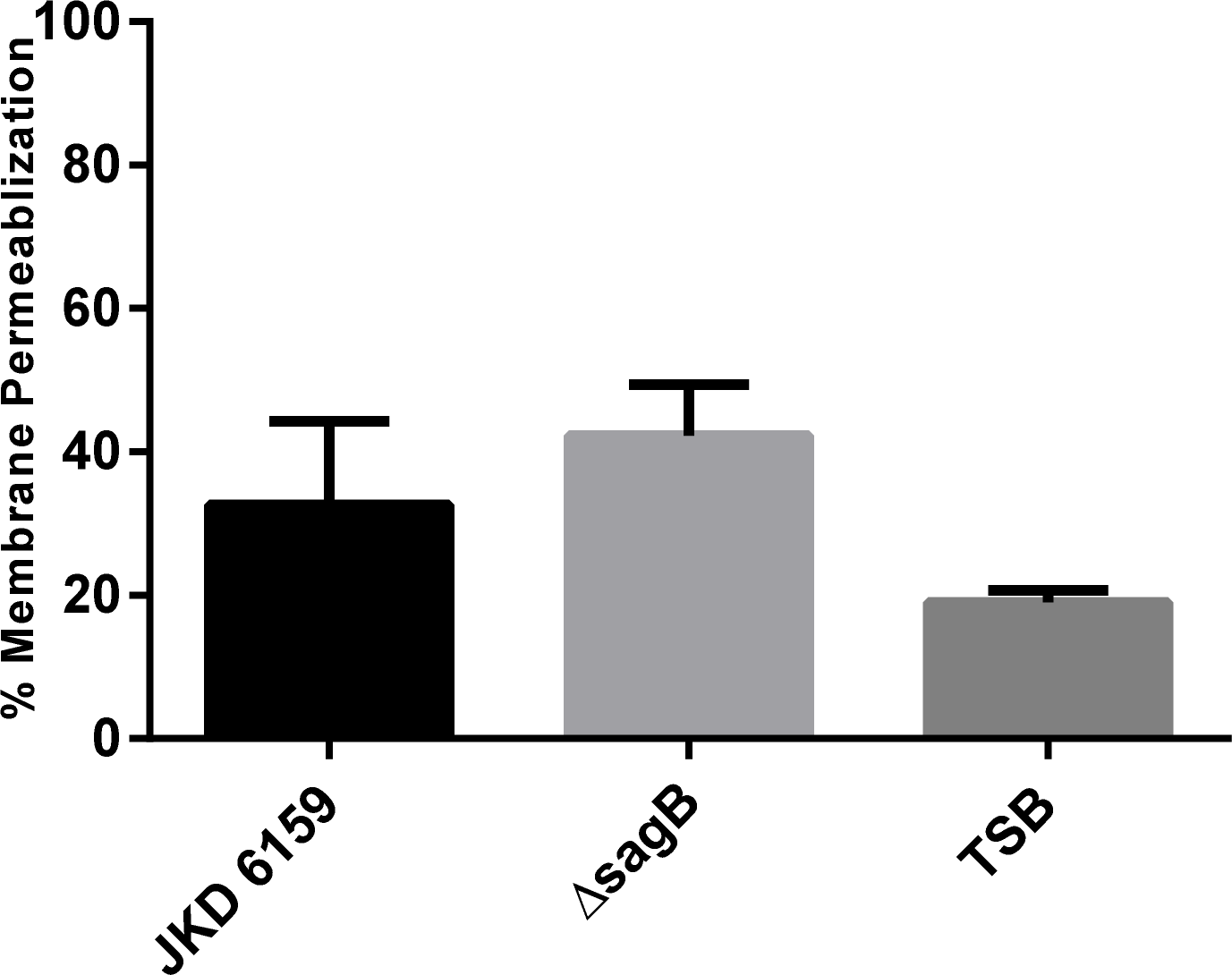
Host cell cytotoxicity from transwell infection of wt JKD6159 and isogenic Δ*sagB* mutant. HaCaT cells were infected at an MOI of 50 for 8 hours and cytotoxicity was assessed using ethidium homodimer assay. Media control was used to assess baseline levels of cytotoxicity.

### Host cell signaling changes

Independent of detectable differences in cytotoxicity, we hypothesized that the putative SLS-like toxin produced by the *sag* cluster in JKD6159 could serve to alter host cell signaling responses in a cell-based infection model. Prior work by our lab has shown that SLS production by GAS can result in a unique cell signaling profile response as measured by signaling array, including an SLS-dependent activation of the proinflammatory transcription factor NF-κB [15]. To assess potential host signaling changes unique to wt JKD6159, we performed an 8 hour transwell infection on keratinocytes (HaCaT cells) at an MOI of 50 of the wt and Δ*sagB* strains. Cells were lysed and an antibody array was carried out (Kinexus). Samples were submitted to Kinexus for the 900P antibody array containing 878 antibodies to various host cell signaling pathway constituents. Signaling changes that were unique to wt JKD6159 as compared to Δ*sagB* included cyclin-dependent protein serine kinases (CDKs) and epidermal growth factor receptor tyrosine kinase (EFGR) (Additional File 2: Table S1).

Changes in EGFR can lead to expression of cytokines from host cells, and previous reports have suggested changes in cytokine expression during *S. aureus* infections can be traced through EGFR activation [21, 22]. We subsequently performed an 80-target immunoblotting array to assess potential differences in cytokine activation between the wtJKD6159 and the isogenic Δ*sagB* strain (abcam). We observed no significant differences in cytokine expression in HaCaT cells infected with wt vs. Δ*sagB* JKD6159 strains (data not shown).

### Antibacterial assays

In select strains of *E. coli*, the microcin B17 peptide is a post-translationally modified peptide similar to SLS that belongs to the larger family of LAPs and acts as a DNA gyrase inhibitor to exert antimicrobial activity [14]. Additionally, LAP family peptides exerting antimicrobial activities have been described in both *Listeria monocytogenes* and *Bacillus amyloliquefaciens* [23, 24]. These LAPs have been shown to target *Staphylococcus aureus* and related *Bacillus* species respectively [23, 24]. Based on these reports, we examined the growth of *E. coli*, GAS, and the common skin commensal *S. epidermidis* when exposed to JKD6159 conditioned media from wt and Δ*sagB* strains. Conditioned media from overnight growth of JKD6159 strains (wt, Δ*sagB*) were sterilized using syringe filtration and added to early growth (OD600 0.2) *E. coli*, GAS, and *S. epidermidis.* We found no significant differences in bacterial growth of each tested bacterial species when treated with wt JKD6159 or the mutants tested (Figure 4).

**Figure 4:**
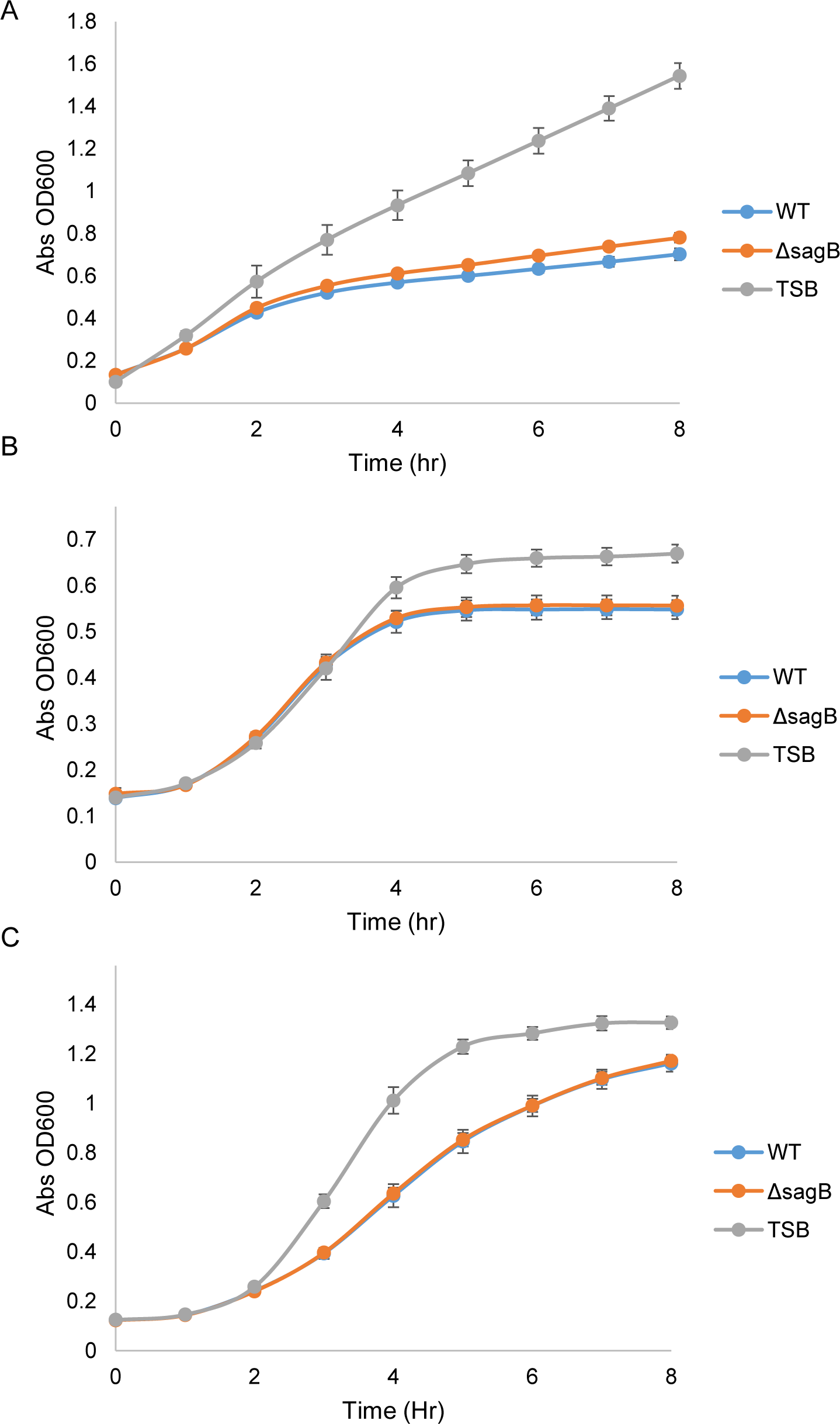
Growth curves of *E. coli, S. epidermidis*, and *S. pyogenes* treated with supernatants from *S. aureus* strain JKD6159 and the isogenic Δ*sagB* mutant. Growths curves of (A) *E.coli*, (B) *S. epidermidis* and (C) *S. pyogenes*. Representative growth curves of triplicate biological replicates.

### Co-culture of GFP *E. coli* with JKD6159 wt and Δ*sagB* strains

In order to further assess any morphological or behavioral changes in *E. coli* when exposed to wt JKD6159, live co-culture of *E. coli* constitutively expressing cytosolic green fluorescent protein (GFP) and JKD6159 wt or Δ*sagB* strains were imaged by epifluorescence microscopy. Stresses introduced to *E. coli* by the *sag* cluster of JKD6159 were hypothesized to be observable *via* live imaging. *E. coli* was diluted to OD600 0.1, and JKD6159 wt or Δ*sagB* strains were added at OD600 0.001. Images acquired over the course of 6 hours revealed no significant morphological differences in *E. coli* between the wt and Δ*sagB* JKD6159 strain co-cultures (Figure 5). Initial evaluation of *E.coli* integrity using wt and Δ*sagB* JKD6159 suggests that this gene cluster does not play a role in bacterial viability of co-cultured strains.

**Figure 5:**
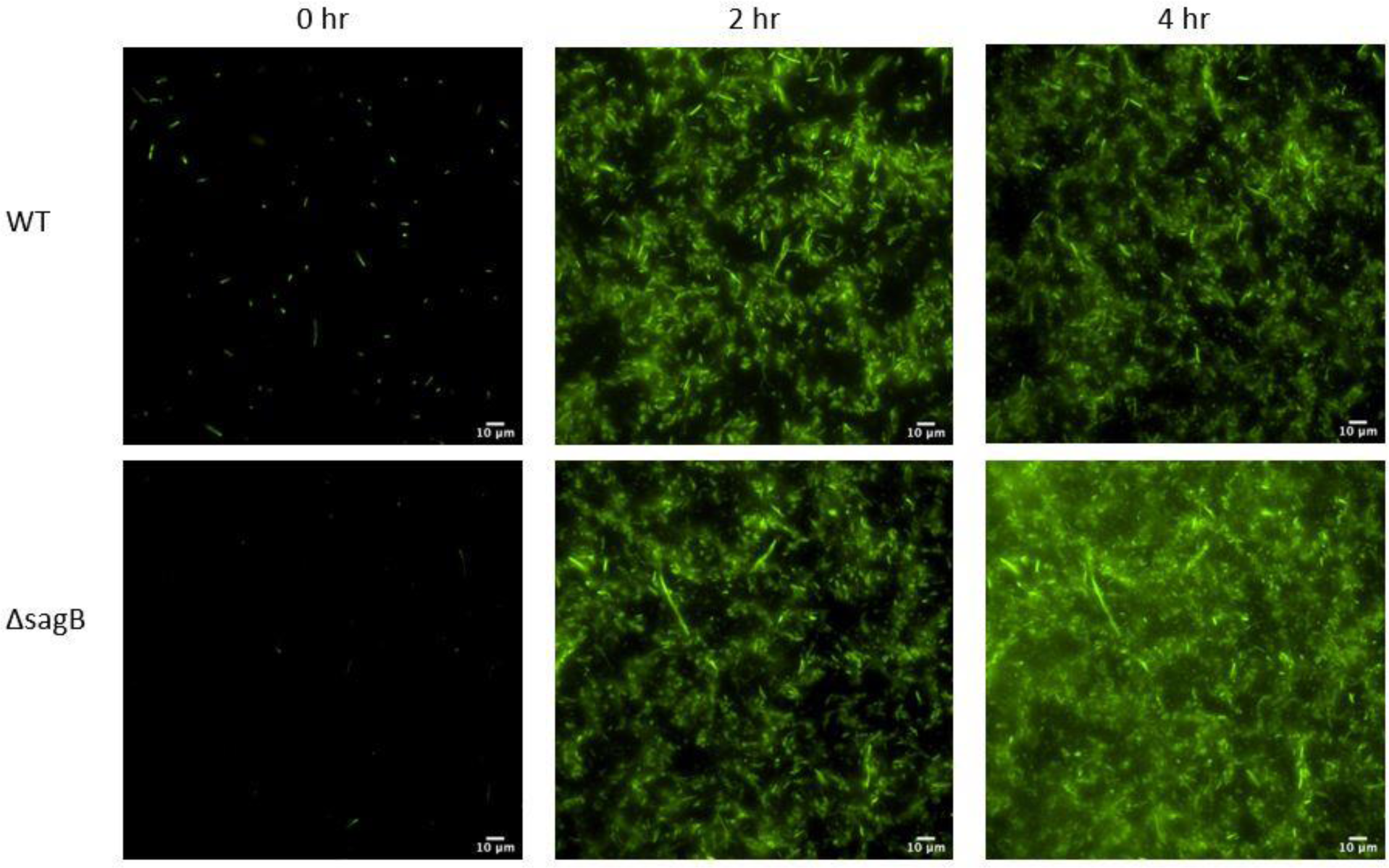
Live imaging co-culture with GFP expressing *E. coli* and JKD6159 wt or Δ*sagB* strain. Select time points of incubation are shown with GFP fluorescence emission.

### Biofilm formation

*S. aureus* biofilms can play an important role in infection, with the ability to create biofilms associated with virulence [25–27]. Bacterial biofilms are associated with a reduction in antibiotic availability, slower clearing infections, and persister cells [26, 28]. To this end we assessed the ability of the wt JKD6159 and Δ*sagB* strains to form biofilms (Figure 6). Overnight cultures were diluted in fresh TSB supplemented with 1% NaCl and 0.5% glucose on PVC coverslips in 6-well plates and allowed to incubate in a humidified environment for 24, 48, or 72 hours. No significant differences in biofilm formation between the strains were observed as measured by crystal violet biofilm assay.

**Figure 6:**
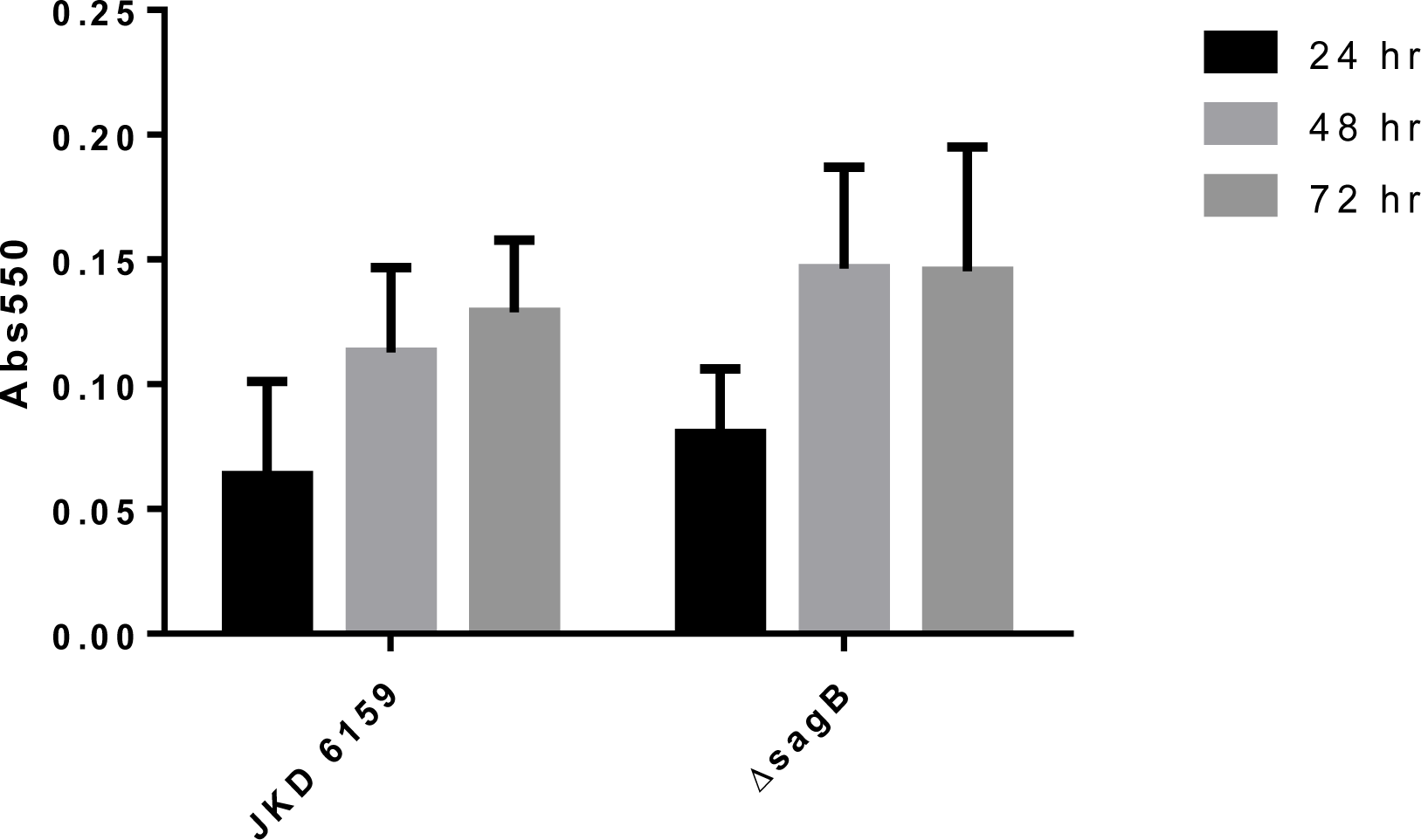
Biofilm formation of the JKD6159 wt and Δ*sagB* strains. Biofilm formation was measured at the indicated time points as measured by crystal violet assay. Results are representative of three biological replicates.

### *In vivo* mouse infections with wt and Δ*sagB* strains

To determine whether the *sag* cluster in JKD6159 contributes to virulence or alters pathogenesis in a manner distinct from the aforementioned *in vitro* virulence assays, *in vivo* mouse infections were carried out according to prior protocols [29, 30]. For five days after intradermal infection with wt JKD6159 and Δ*sagB* strains, mice were weighed and then sacrificed on day 5. Lesions at the infection site were measured and excised for CFU enumeration (Figure 7). The outcomes measured indicated no alterations in the course of infection as a result of the absence or presence of the *sag* cluster in JKD6159.

**Figure 7:**
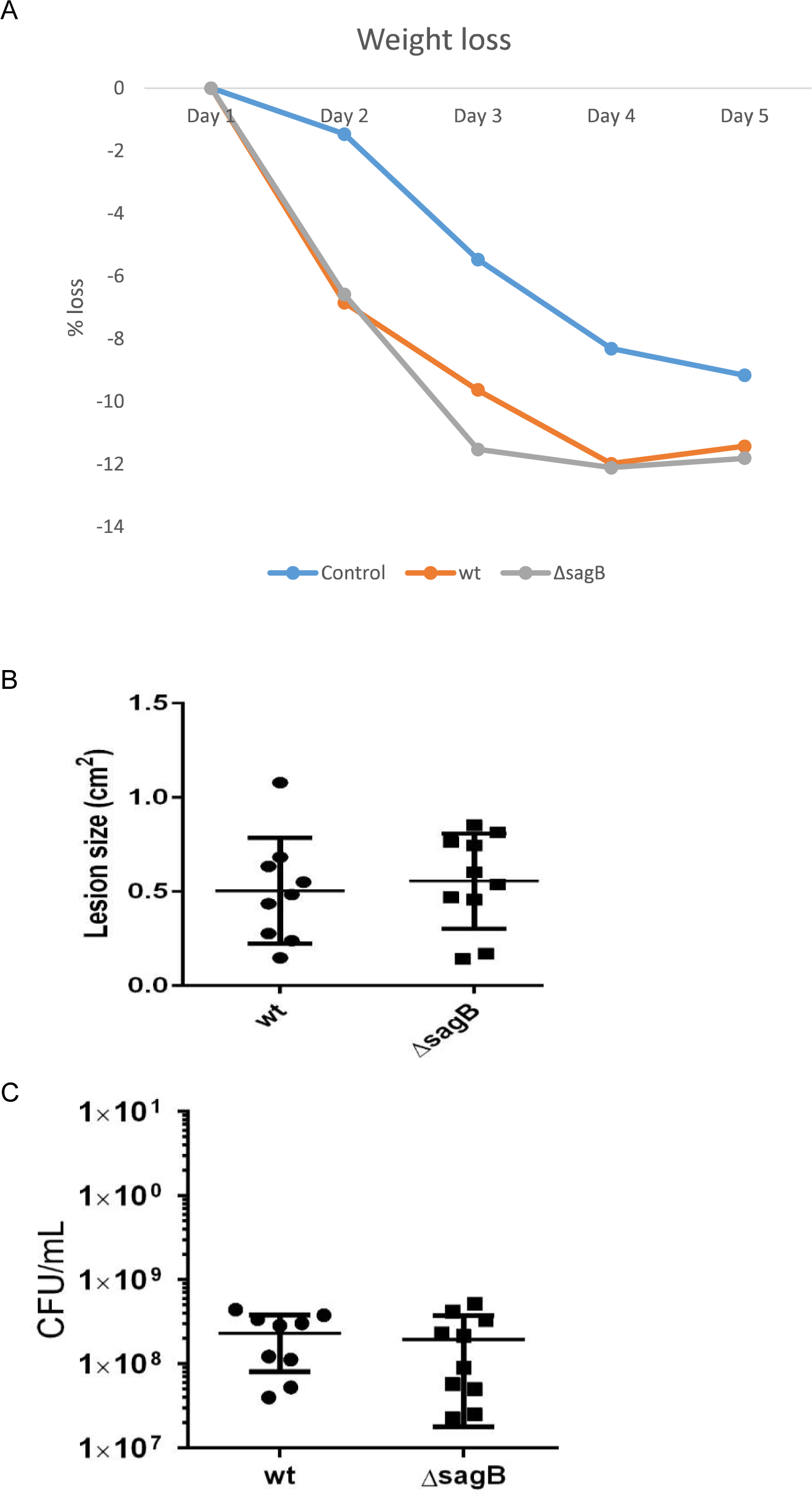
*In vivo* infection. (A) Per cent weight loss during the course of infection with each of the JKD6159 wt and Δ*sagB* strains as indicated. (B) Lesion size following subcutaneous infection on day 5 with JKD6159 wt and Δ*sagB* strains. (C) CFU/mL recovered from the lesion excised 5 days following subcutaneous infection with JKD6159 wt and Δ*sagB* strains.

### 2D proteomics to assess differential protein expression between wt and Δ*sagB* JKD6159

In order to determine if disruption of the *sag* cluster in the JKD6159 strain of *S. aureus* plays a role in the regulation of specific JKD6159 bacterial proteins, we undertook a 2D proteomic approach to visualize the proteome. After separation of JKD6159 lysates by size and isoelectric point, spots that were consistently different between wt and Δ*sagB* mutant strain in duplicate technical and biological replicates were excised and identified using mass spectrometry (Additional Files 3, 4: Figure S2, Table S2).

Proteins that were more abundant in the wt JKD6159 strain compared to the Δ*sagB* strain included threonine synthase, 2 oxoisovalerate dehydrogenase subunit alpha, carbamate kinase, NH3 dependent NAD synthetase, succinate CoA ligase ADP forming alpha subunit, thioredoxin disulfide reductase, Putative glucokinase ROK family, and D isomer specific 2 hydroxyacid dehydrogenase family protein. Proteins that were more strongly expressed in the Δ*sagB* strain included uroporphyrinogen decarboxylase, FolD, Global transcriptional regulator catabolite control protein A, hydroxymethylbilane synthase, FAD dependent pyridine nucleotide disulphide oxidoreductase, and S adenosyl methyltransferase MraW.

Threonine synthase is responsible for the conversion of L-phosphohomoserine to Lthreonine via the addition of water [31]. 2 oxoisovalerate dehydrogenase subunit alpha is part of a large protein complex in the TCA cycle, and is responsible for the conversion of alpha-keto acids to acetyl-CoA [32]. Carbamate kinase is responsible for transferring a phosphate from carbamoyl phosphate to ADP thereby generating ATP [33]. NH3 dependent NAD synthetase is responsible for the final step in generating NAD+ from deamindo-NAD+ [34]. Succinate CoA ligase ADP forming alpha subunit is involved in the substrate level phosphorylation of ADP in the TCA cycle [35]. Thioredoxin disulfide reductase catalyzes the reaction of NADP+ to NADPH [36]. Putative glucokinase ROK family plays a role in the glucose metabolism pathway [37]. D isomer specific 2 hydroxyacid dehydrogenase family protein is a member of the carbohydrate metabolism pathway and it converts keto acids into chiral hydroxy acids [38].

The proteins that are highly expressed in the Δ*sagB* strain of JKD6159 are as follows: Uroporphyrinogen decarboxylase catalyzes the first step of the tetrapyrrole pathway [39]. FolD is a bifunctional enzyme that generates NADPH and 10formyltetrahydrofolate [40]. Global transcriptional regulator catabolite control protein A (CcpA) is a master regulator involved in many catabolic pathways [41].

Hydroxymethylbilane synthase is involved in polymerization of porphobilinogen into 1hydroxymethylbilane [42]. FAD dependent pyridine nucleotide disulphide oxidoreductase is responsible for catalyzing disulfide bond formation [43]. S-adenosyl methyltransferase MraW is a methyltransferase that methylates 16s rRNA [44].

### Evolutionary dynamics of the *sag* cluster in *S. aureus*

In order to investigate the evolutionary origin and dynamics of *sagB* and the *sag* cluster in *S. aureus*, we identified all strains of *S. aureus* and other *Staphylococcus* species containing the *sag* cluster (Figure 8A). The *sag* cluster was only found in a small proportion of known strains of *S. aureus* (49 out of 7999) and is located at a site between the gene *gnd* (phosphogluconate dehydrogenase) and gloxalase (Figure 8B). Among the genus of *Staphylococcus*, the *sag* cluster was additionally found to be present in *Staphylococcus lugdunensis* (*S. lugdunensis*) and two other *Staphylococcus* species, *i.e.*, *Staphylococcus argenteus* (*S. argenteus*) and *Staphylococcus schweitzeri* (*S. schweitzeri*) (Figure 8A). The latter two species are closely related with *S. aureus* and were just recently reclassified as separate lineages from *S. aureus* [45, 46]. The *sag* cluster was also identified in two strains of *S. cohnii*. Based on the distribution and sequence properties of the *sag* cluster in the genus of *Staphylococcus*, it is tempting to postulate that an ancestral strain of *S. aureus* may have acquired the *sag* cluster from *S. lugdunensis* or their common ancestor by horizontal gene transfer and thereafter suffered from genetic loss of the *sag* cluster based on three lines of evidence. First, the similarity of the *sag* cluster between *S. lugdunenesis* and *S. aureus* is averaged to 94%, higher than the average genome-wide similarity of 80% between the two species. Second, the *sag* cluster is located at the same loci in the genomes of different strains of *S. aureus*, indicating the clonal inheritance of the *sag* cluster from an early ancestor. Third, the *sag* cluster was identified in all 19 known strains of *S. lugdunensis*, but only identified in a small proportion of *S. aureus* strains (49 out of 7999) and *S. argenteus* strains (17 out of 111) with known genomic information. It should also be noted that the evolutionary dynamics of the *sag* cluster remains to be further determined as more genomic information from *S. lugdunensis* becomes available.

**Figure 8:**
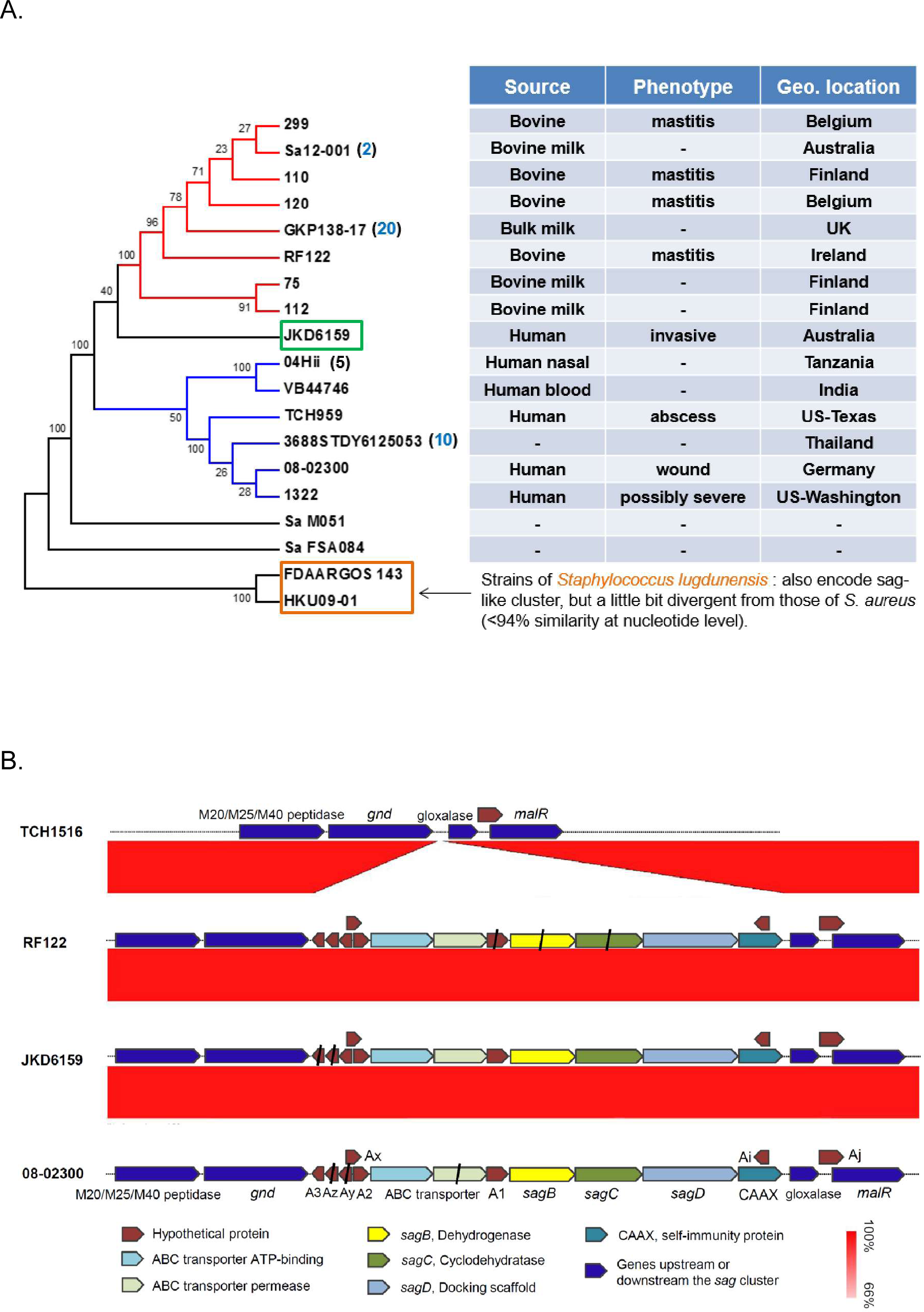
Other *S. aureus* strains containing the sag-like cluster and their phenotypes.

(A) *S. aureus* strains from bovine sources and human sources are separately clustered (red and blue). However, the human invasive isolate JKD6159 is observed to be distantly clustered with other human isolates (green box). S. lugdunensis strains containing a sag-like gene cluster are also included in phylogenetic tree (brown box).
(B) Organization of *S. aureus* strains containing the sag-like cluster. *S. aureus* strains TCH1516, RF122, JKD6159 and 08-2300 harbor a sag-like gene cluster in similar insertions locations and have similar organization of genes to each other.

We further propose that the loss of *sag* cluster among the large *S. aureus* population is driven by neutral or adaptive selection. The absence of the *sag* cluster in most of the known strains of *S. aureus* and the truncation of multiple genes within the *sag* cluster among the *sag* cluster-containing strains of *S. aureus* indicate that the *sag* cluster might be under neutral selection and dispensable for hemolysis and human virulence of *S. aureus*, as demonstrated in the current study. *S. aureus* produces multiple cytolytic toxins, which work synergistically in causing membrane damage and are required in various host diseases. Among them, the α-hemolysin has been shown to be a major cytolysin and key virulence factor of *S. aureus* in causing skin/soft tissue infections and invasive diseases [47]. The γ-hemolysin was reported to act in concert with α-hemolysin to give rise to septic arthritis in a murine model [48]. The β-hemolysin has shown to be less virulent for general cell damage [48], and mainly responsible for increasing the sensitivity of red blood cells to other toxins [49, 50]. We found that the γ-hemolysin and β-hemolysin are highly conserved among the genomes of *S. aureus*, in contrast with the sparse distribution of the *sag* cluster. It is probable that the cytolytic toxins and other virulence genes encoded by *S. aureus* have been sufficient for its pathogenesis in a wide range of conditions and the acquisition of *sag* cluster by the ancestral strains might be an accidental event during its evolution. The hypothesis of neutral selection of the loss of *sag* cluster also implicates population bottlenecks preceding the global expansion of the clones with deleted *sag* cluster [51], which is not uncommon for pathogenic bacteria in colonizing environments with antibiotic treatments [52].

## Discussion

We have utilized various approaches to study the function of a *sag* gene cluster found in the CA-MRSA strain JKD6159. As the specific protoxin gene was not readily apparent, we engineered a *sagB* deletion in the cluster, and prior work has shown that deletion of any of the 9 genes found in the cluster led to loss of function of the active toxin [9]. The peptides that are in the LAP family are responsible for a range of phenotypes, including antibiotic activity, host cytotoxicity, and hemolysis [10, 14, 15]. Based on the range of activities for which the *sag* cluster in JKD6159 could potentially be responsible, we undertook a comprehensive panel of host and bacterial phenotype testing to potentially identify a role of the *sag*-like gene product in JKD6159. With respect to host cell pathogenicity, we showed that the *sag* cluster has no effect on hemolysis or cytotoxicity and minimal effects on host signaling pathways. Similarly, we observed no apparent inhibition on the growth of *E. coli, S. epidermidis*, or GAS when treated with conditioned media from wt JKD6159.

Our comparative proteomic analysis of wt vs. *sagB* mutants revealed changes in levels of several metabolic proteins, possibly owing to perturbed metabolic flux introduced by the enzymatic activity of the *sag* cluster or production of the SLS-like peptide product. The oxidative dehydrogenation of substrate peptides carried out by SagB is thought to be flavin mononucleotide (FMN)-dependent, and the activity of SagB could deplete the pool of cellular FMN enough to trigger regulatory feedback signals that lead to the observed differences in metabolic protein abundances. While the impact of many of these metabolic pathways is unclear in regards to the *sagB* deletion in JKD6159, the regulatory protein CcpA has been implicated in the virulence of both GAS and *Streptococcus pneumoniae* (*S. pneumoniae*) and was found in higher abundance in Δ*sagB* JKD6159 [41, 53]. In both species, deletion of *ccpA* results in decreased virulence. Deletion of *ccpA* in *S. pneumoniae* results in decreased colonization of mouse lungs [41]. In GAS the loss of CcpA results in reduced infectivity in mouse models of infection and reduced colonization. Interestingly, purified CcpA binds to the promoter region of [53]. It is possible that the increase in CcpA abundance in the Δ*sagB* strain is compensatory for *sag* cluster deficiency or that the putative LAP product of the *sag* cluster exerts a negative feedback on CcpA. Further elucidating the interplay between the *sag* cluster and CcpA could shed light on the role of the *sag* cluster in JKD6159.

Recent work by Quereda et al. has shown that the toxin Listeriolysin S (LLS), the LAP product of a *sag* cluster found in *Listeria monocytogenes* (*L. monocytogenes*), only affects the microbiome during gut infections in mice [24]. Similar to our observations, the authors found no or minimal differences in the hallmark hemolysis, cytokine signaling, cytotoxicity, or virulence typical of SLS that could be attributed to LLS [24]. Yet during gut infections, the mouse microbiome is altered between the wt and LLS-deficient strains of *L. monocytogenes* via selective inhibition of a handful of bacterial species within the complex polymicrobial community of the gut [24]. Moreover, LLS expression was highly context-dependent within the host gastrointestinal tract, and specific antimicrobial activity could be demonstrated *in vitro* only with a strain of *L. monocytogenes* constitutively expressing *llsA*, the LLS precursor peptide.

These data suggest that complete assessment of the biological impact of the *sag* cluster in *S. aureus* JKD6159 may require analysis under appropriate context within the host, extensive genetic regulation and expression analysis, and genetic constructs that bypass host context constraints to allow *ex vivo* studies on function and mechanisms. The *sag* cluster may thus have distinct effects on the host and the surrounding polymicrobial community only when under specific conditions relevant to human infection, and direct antimicrobial activity may be selectively exclusive of the strains of GAS, *S. epidermidis*, and *E. coli* examined here. With respect to the rising threat of MRSA, it is critical to proactively examine the impact of acquisition of novel biosynthetic clusters like the *sag* cluster of JKD6159 to guide clinical decision making. To that end, the functional breadth of the handful of LAPs studied thus far and complex regulation of biological functionality demands further rigorous study of LAP biosynthetic clusters and their effects, particularly in the context of established human pathogens.

## Conclusions

Using bioinformatic tools, we identified a Streptolysin S (SLS) like biosynthetic gene cluster in the methicillin resistant *S. aureus* (MRSA) isolate JKD6159, and generated a knockout mutant of *sagB* in the biosynthetic gene cluster. We found no differences in hemolysis, cytokine signaling, host cell cytotoxicity, or virulence in mice between our wt and Δ*sagB* mutant. There were also no differences in growth inhibition of other bacteria or in *S. aureus* biofilm formation. Analysis of the evolutionary dynamics of this sag-like cluster show its absence in a large proportion of *S. aureus* strains and suggests it may be under neutral selection. Our data suggests that further work is necessary to determine the biological function of this sag-like cluster which may require highly specific host contexts.

## Abbreviations

GAS: Group A *Streptococcus*
SLS: Streptolysin S
*sag*: Streptolysin S-associated gene
MRSA: Methicillin resistant *Staphylococcus aureus*
CA-MRSA: Community acquired MRSA
HA-MRSA: Hospital acquired MRSA
LAPS: Linear azole containing peptides
OD: Optical density
CFU: Colony forming units
LB: Lysogeny broth
TSB: Tryptic soy broth
PBS: Phosphate buffered saline
MOI: Multiplicity of infection
PVC: Polyvinyl chloride
PCR: Polymerase chain reaction
GFP: Green fluorescent protein
ABC: Ammonium bicarbonate
FBS: Fetal bovine serum
DMEM: Dulbecco’s modified eagle medium
TBST: Tris buffered saline with tween
ECL: Enhanced chemiluminescence
IPG: Immobilized pH gradient
DTT: Dithiothreitol
HRP: Horseradish peroxidase
Cm: Chloramphenicol
BHI: Brain heart infusion
ATC: Anhydrotetracycline

## Methods

### Bacterial strain information

*S. aureus* strain JKD6159 was the generous gift of Dr. Timothy Stinear (University of Melbourne). Vector pIMAY and pCL55 were generous gifts of Dr. Ian Monk and Dr. Taeok Bae (Indiana University). *E. coli* strain IM93B was a generous gift of Dr. Ian Monk (University of Melbourne) and Belgian Coordinated Collections of Microorganisms (BCCM). Unless otherwise mentioned all strains were grown at 37°C.

### Preparation of competent bacteria

*E. coli* strain IM93B was prepared as described previously [54, 55]. JKD6159 was grown in 3 mL tryptic soy broth (TSB) overnight and then diluted in 300 mL TSB. Cultures were allowed to grow to an optical density (λ = 600 nm; OD600) of 0.5. JKD6159 was then centrifuged 4°C at 3500 RPM. The bacteria were washed once using 60 mL ice-cold 0.5 M sucrose H2O, then washed three more times in 9 mL 0.5M cold sucrose H2O. Cells were then resuspended in 3 mL 0.5M ice-cold sucrose H2O, aliquoted into 100 µL, and snap frozen in an ethanol dry ice bath and stored at −80°C prior to use.

### Generation of *sagB* deletion mutant in JKD6159

Isogenic gene deletions of *sagB* were obtained following the protocols in [54] and [56]. Primer sequences for the generation of *sagB* mutants are found in supplementary Table S3 (Additional File 5). *≈*800 bp regions of the chromosome upstream and downstream of the *sagB* gene to be deleted were amplified using sagB KO downF, sagB KO downR, sagB KO upF, and sagB KO upR as found in Table S3. PCR reaction products were visualized on a 1% agarose gel (Sigma-Aldrich) containing a 1:10,000 dilution of GelRed (Biotium). pIMAY and the upstream region were digested with EcoR1 and Not1 (New England BioLabs) according to manufacturer instructions. Digested DNA was then purified using QIAquick PCR purification kit from Qiagen according to manufacturer instructions. The digested upstream region and pIMAY were ligated together using T4 DNA Ligase (New England BioLabs) according to manufacturer instructions. 1 µL ligation reaction was then transformed into *E. coli* strain One Shot ®Top10 (Invitrogen) according to manufacturer instructions and plated on LB containing 10 µg/mL chloramphenicol (Cm 10, SigmaAldrich) for selection. *E. coli* clones able to grow on Cm 10 were then PCR amplified for the presence of the upstream region using primer set T7F and T7term. Phusion High-Fidelity DNA polymerase was used according to manufacturer instructions for reactions using a 2-minute initial denature at 98°C, followed by 30 cycles of 30 s at 96°C, 30 s at 55°C, and 30 s at 72°C, and finally held at 10°C (New England Biolabs). This was the standard PCR reaction used unless otherwise noted. Clones that contained the insert as shown on an agarose gel were then grown in 10 mL LB (Fisher Scientific) Cm 10 and subjected to Zippy plasmid Miniprep Kit (Zymo Research) according to manufacturer recommendations. Purified plasmid was then quantified on a NanoDrop 2000 (Thermo Scientific). 300 ng of purified plasmid was then sequenced using T7F and T7term on an Applied Biosystems 96-capillary 3730xl DNA Analyzer in the Genomics and Bioinformatics Core Facility of the University of Notre Dame. Sequencing results were analyzed using Sequencher (Gene Codes). Plasmids containing the correct insert were then digested along with the downstream region using Kpn1 and EcoR1 (New England BioLabs), and digested DNA was purified with QIAquick kit (Qiagen) according to manufacturer recommendations. DNA was then ligated using T4 DNA ligase (New England BioLabs) and 1 µL of the ligation reaction was transformed into *E. coli* strain One Shot ®Top10 (Invitrogen). Clones were selected for on LB (Fisher Scientific) Cm 10 µg agar plates, and individual colonies were checked with PCR for bands corresponding to ∼1600 base pairs indicating that the vector contained both the upstream and downstream region. PCR was repeated as above with the only change being the extension time was increased to 1 minute from 30 seconds. Positive clones were grown further and miniprep was performed to isolate plasmids, and sequenced as above. Plasmids that contained the correct upstream and downstream region as shown by Sanger sequencing was then transformed into *E. coli* strain IM93B. Competent IM93B was thawed on ice, and then added to chilled 0.1 cm electroporation cuvettes (BioRad). 100 ng of pIMAY containing the upstream and downstream regions to *sagB* was added to the competent IM93B and electroporation was carried out at 100 Ω, 1.8 kV, 25 µF using a BioRad Gene Pulser Xcell. 500 µL SOC media (Invitrogen) was added, and the bacteria were allowed to recover shaking at 37°C for 45 minutes. Cells were then plated on LB Cm 10 µg and allowed to grow overnight. Colonies were checked for presence of pIMAY with correct inserts using PCR, miniprep, and sequencing as above to ensure that no additional mutations had occurred. Clones containing the correct insert were then grown up for plasmid isolation using a Qiagen QIA Filter Plasmid Maxi kit according to instructions. Competent JKD6159 was thawed on ice for 15 minutes then spun at 8000 g for 3 minutes in an Eppendorf 5415R centrifuge. Bacteria were then resuspended in 100 µL 10% glycerol containing 0.5M sucrose (Fisher Scientific). 2 µg plasmid passed from IM93B was added to the competent JKD6159. Plasmids were introduced into the JKD6159 bacteria via electroporation in a 0.1 cm electroporation cuvette (BioRad) at 100 Ω, 2.5 kV, 25 µF. After electroporation the JKD6159 was allowed to recover in 1 mL Brain Heart Infusion (BHI, Fluka) for one hour at 37°C. Cells were centrifuged at 8000 g for 3 minutes using the Eppendorf 5415R and resuspended in 100 µL BHI and plated on BHI Cm 10 µg/mL agar plates and incubated at 28°C for 2 days. Colonies that grew under selective pressure at 28°C were homogenized in 200 µL TSB, diluted 100 fold, and 100 µL were plated on BHI Cm 10 µg/mL at 37°C to force the first integration of the pIMAY plasmid. Colonies that grew were grown in 1 mL BHI Cm 10 µg/mL and genome DNA recovered using MasterPure™ Gram Positive DNA Purification Kit from Epicenter. Integration of the plasmid was verified using pIMAYF and BR KO chromosome (Table S3). Clones that successfully integrated the vector were grown overnight in BHI at 28°C. Dilutions of 10^−6^ and 10^−7^ were plated on BHI agar containing 1 µg/mL anhydrotetracycline (ATC, Sigma-Aldrich) at 28°C for 24-48 hours. Large colonies were patched onto both BHI ATC 1µg/mL and BHI Cm 10 µg/mL and allowed to grow overnight at 37°C. Colonies that were able to grow on BHI ATC but not on BHI Cm were selected as potential positive candidates for the gene deletion. Cm sensitive colonies were grown in BHI overnight at 37°C overnight and genome DNA purified using the MasterPure™ Gram Positive DNA Purification Kit. Chromosomal primers BF KO chromosome and BR KO chromosome were used to verify the loss of the *sagB* gene. PCR for the verification of gene deletion is the same as above however the extension time was increased to two minutes. 50 mL of overnight culture of the *ΔsagB* were grown in BHI, centrifuged at 2500 g for 10 minutes, resuspended in 1 mL BHI with 30% glycerol (Fisher Scientific) and stored at −80°C.

### Sheep blood hemolysis assay

1 mL defibrinated sheep blood (Thermo Fisher Scientific) was washed in 40 mL 4°C PBS to remove the free heme from the lysed red blood cells (RBC). RBCs were then centrifuged at 4°C 1200 RPM for 10 minutes. RBCs were determined to be free of lysed cells when the supernatant was mostly clear, usually two wash cycles. Intact RBCs were resuspended in ice cold PBS 40 mL, and aliquoted into 1.5 mL snap top tubes. Sterile conditioned media was generated from the wt and Δ*sagB* JKD6159 strains by growing the bacteria overnight in TSB media, centrifuging the bacteria at 3500 RPM and sterile filtering the supernatant using Millipore 0.22 µM luer lok syringe filters. This conditioned media was then added to the washed RBCs to a ratio of 1:3. The reactions were then incubated at 37°C for the designated time points. To read hemolysis, the tubes were centrifuged at 1200 RPM centrifuge for 10 minutes. 100 µL of the supernatant was transferred in Eppendorf UVette cuvettes before being read at O.D. 450 nm to measure free heme. Hemolytic action was calculated by normalizing the lysis to 0.01% Trition X-100 positive control and TSB negative control. Experiments were conducted in triplicate.

### Antibacterial assays

Overnight cultures of *E. coli, S. pyogenes* and *S. epidermidis* in LB media were diluted in LB (*E. coli*), TH (*S. pyogenes*), or Nutrient Broth (*S. epidermidis*) to an OD600 of ∼0.1. Conditioned media from the JKD6159 wt and Δ*sagB* strains were prepared by overnight growth in TSB media followed by centrifugation at 3500 RPM and sterile filtration through Millipore 0.22 µm luer lok syringe filter. Conditioned media was added to the diluted overnight cultures of *S. epidermidis, S. pyogenes* and *E. coli* at 1:3 ration in Eppendorf 96 well plates. The plates were read on a Synergy H1 microplate reader by BioTek. The plate reader was heated to 37°C, set to continual shaking, and read OD600 every 10 minutes for 16-24 hours. Experiments were conducted in biological and technical triplicate.

### Live imaging of JKD6159 and *E. coli*

Co-culture experiments were also carried out using GFP *E. coli* pND1308 (pCRFcr/GFP) TOP10 (Gift of Zhong Liang). *E. coli* and JKD6159 were grown overnight before being diluted to OD600 0.1 (*E. coli*). Wt and Δ*sagB* JKD6159 was diluted 1000 fold less than *E. coli* before being added to glass bottom petri dish (MatTek) containing 1 mL of 1:1 LB:TSB. Live imaging was acquired on Nikon Eclipse Ti-E Inverted Microscope on a 60X oil immersion NA1.40 objective. Images were captured using EMCCD camera (Andor Ixon Ultra 88 ECCD Oxford Instruments) with a 10 ms exposure time. Petri dish was placed in an environmental chamber at 37°C for 12 hours and imaged continuously for 12 hours in DIC and FITC channels.

### Biofilm formation assay

Overnight cultures of JKD6159 wt and Δ*sagB* were grown in TSB. Cultures were then diluted 1:100 in TSB supplemented with 1% NaCl and 0.5% glucose. 2 mL of diluted culture was pipetted on top of sterile PVC coverslips (Fisher Scientific) in 6-well plates and placed in a humidified 37°C, 5% CO2 incubator for 24, 48, or 72 hours. Following the time point, coverslips were washed 2x with ddH2O, and 2 mL of 0.1% crystal violet was added and incubated at room temperature for 10 minutes. Crystal violet was then aspirated, and wells were washed 2x with ddH2O. Coverslips were allowed to air dry, and remained at room temperature until solubilization with 33% acetic acid in water and reading at OD on a Synergy H1 microplate reader by BioTek. Experiments were conducted in technical triplicate and biological triplicate.

### 2D proteomics of JKD6159 wt and Δ*sagB*

The procedure below was adapted from [57, 58]. wt JKD6159 and Δ*sagB* JKD6159 were grown in 10 mL TSB at 37°C for 48 hours. Bacteria were centrifuged at 4°C at 2500 g for 10 minutes. The pellet was then washed twice in 500 µL PBS containing SigmaFast protease inhibitor tablet (Sigma). Cells were resuspended in 500 µL PBS with protease inhibitor and added to ∼200 µL 0.1mm glass beads before being placed in Mini Bead Beater (Biospec) for 4 cycles of 30 seconds each, with one minute on ice between each cycle. Upon conclusion of mechanical disruption, 500 µL of 2D protein Extraction Buffer V (GE Healthcare), 6 µL Destreak reagent (GE Healthcare), and 2.5 µL IPG buffer 3-10 pH NL (GE Healthcare) was added to samples, and samples were allowed to incubate at room temperature for one hour with vortexing every 10 minutes for 10 seconds. Lysates were then centrifuged at 4°C at 16,000 g for 30 minutes. Supernatants were then collected and centrifuged at 50,000 g for 1 hour, twice, to remove any additional insoluble material. Supernatant containing the soluble proteins was quantified using Comassie Plus Assay Kit (Thermo Scientific) according to manufacturer instructions. 300 µg total protein was prepared for 2D proteomics using the 2D Clean Up Kit (GE Healthcare) according to manufacture instructions. Proteins were then resuspended in 250 µL Destreak Rehydration Solution (GE Healthcare), 6 µL of DTT 1M, and 6 µL of IPG 3-10 pH NL (GE Healthcare). Samples were vortexed for 15 minutes, and any remaining insoluble material was removed via centrifugation at 16000 g for 1 minute. Samples were then allowed to adsorb into Immobiline DryStrip 3-10 pH NL 13 cm (GE Healthcare) overnight. The first dimension separation was achieved using an Ettan IPGphore 3 (GE Healthcare) following suggested protocols. Prior to second dimension separation, the strips were equilibrated by incubation at room temperature in 5 mL SDS equilibration buffer (50mM Tris-HCL, 6M urea, 30% glycerol, 2% SDS, 0.01% bromophenol blue) supplemented with 650 µL of 1M DTT for 15 minutes, followed by 5 mL SDS equilibration buffer supplemented with 1350 µL of 1M iodoacetamide. Second dimension separation was made on a 12.5% polyacrylamide gel using the SE600 Ruby system (GE Healthcare). Mark 12 unstained standard was used for molecular weight estimations. The DryStrip was covered with sealing agarose (BioRad). The gel was then run at 600 V, 30 mA, 100 W for 15 mins, then 600 V, 60 mA, 100 W for 4-6 hours. mA was set as the limit for the power supply. Gels were then stained using Sypro Ruby according to manufacturer instructions and imaged using a Typhoon imager FLA9500 (GE Healthcare). Data was analyzed using Delta 2D (Decodon). Spots that were determined to be significantly different were manually excised and subjected to mass spectrometry.

Gel spots were excised from the gel, destained in 25mM ammonium bicarbonate (ABC) in 50% water 50% acetonitrile. The spots were then placed in an Eppendorf Speed Vac at 35°C until spots were completely dried. 25 µL of 10 mM DTT in 25 mM ABC was added to dried spots and incubated at 56°C for 1 hour. The supernatant was then removed and spots were washed 2x by adding 100 µL 25 mM ABC and vortexed for 10 minutes at room temperature. Gels were dehydrated using 25 mM ABC in 50% acetonitrile 50% water and vortexed for 5 minutes. Solution was then removed and new 25 mM ABC in 50% acetonitrile 50% water was added again and vortexed an additional 5 minutes. Gel spots were again placed in an Eppendorf Speed Vac at 35°C until completely dried. 500 ng of Trypsin Gold (Promega) in 25mM ABC was added to gel spots and allowed to rehydrate on ice for 30 minutes. If needed, 25 mM ABC was added to cover the gel spots after the rehydration. Trypsin was allowed to digest at 37°C overnight. Supernatant containing peptides was reserved and the spot was covered in extraction buffer (45% water, 50% acetonitrile, 5% formic acid (Fisher Optima) and vortexed for 30 minutes. Extraction of peptides from spot was repeated once and all supernatants were pooled and concentrated using a Speed Vac adjusted to 10µL total volume. Peptides were then run on a Thermo Fisher Q-Exactive mass spectrometer at Notre Dame proteomics core facility. The 2D proteomic studies were carried out biological and technical duplicate.

### Cytotoxicity and host cell signaling changes between wt and Δ*sagB* JKD6159

HaCaT keratinocyte cells were used for all cytotoxicity and cell signaling assays. Unless otherwise indicated, cells were kept in a humidified, 37°C, 5% CO2 environment in DMEM (Gibco) containing 10% fetal bovine serum (Biowest) in T75 flasks. In cytotoxicity assays, HaCaT cells were grown in DMEM lacking FBS in 24 well plates. HaCaTs were allowed to grow to 80-90% confluency. Prior to infection, cells were washed twice with sterile PBS (Gibco) and then fresh DMEM media was added to the wells. wt and Δ*sagB* JKD6159 bacteria were added at an MOI of 50 bacteria/1 host cell in a Millipore Transwell 0.4 µm. Cells and bacteria were allowed to incubate for 8 hours at 37°C, 5% CO2. Upon completion of the time points, the transwells were removed, and HaCaTs rinsed with PBS. Ethidium homodimer (Sigma Aldrich) at a concentration of 4 µM was added, and the cells were allowed to incubate at room temperature in the dark for 30 minutes. The plate was then read on a BioTek Synergy H1 at excitation 528 nm, and emission 617 nm to obtain a treated level of fluorescence. Saponin (Sigma) was added at a concentration of 0.1% w/v to lyse the remaining cells and florescence reading was repeated to allow calculation of cytotoxicity.

Cell signaling data was obtained by using a Kinexus KAM-900P antibody array. HaCaTs were treated as above for the cytotoxicity assay, but upon completion of the 8 hour time point, cells were washed twice in ice cold PBS, before being lysed using Kinexus lysis buffer plus protease inhibitor. 8 wells of the 24 well plate were lysed for each of the wt and Δ*sagB* conditions. The cells were sonicated for 4×10 seconds with 30 seconds on ice in between each pulse. The lysates were then spun at 16,000 g at 4°C for 30 minutes to remove cell debris. Lysates were then stored at −80°C until they could be sent to Kinexus for antibody array analysis.

Cytokine antibody array analysis was also performed using the Abcam Human Cytokine array kit with 80 targets. Prior to transwell infection as outlined above, the HaCaT cells were serum starved for 16 hours before being infected by the wt and Δ*sagB* mutant strain for eight hours. Infections were carried out in triplicate wells and 400 µL from each well was collected and pooled before being frozen at −20°C until the array analysis was performed. The antibody array was performed as instructed by the manufacturer, with the membranes being incubated overnight at 4°C with the supernatants collected from the HaCaTs, and overnight incubation at 4°C with the primary antibodies. Membranes were imaged using an Azure Biosystems c600 imager, and images were then quantified using ImageJ.

Cytokine hits IL-10, I-309, osteopontin, and MIP-1δ were verified via western blots. IL-10 antibody was from Abcam number ab34843, osteopontin was Abcam number ab8448, MIP-1δ was R&D systems AF628, and I-309 was from Abcam ab109788. Supernatants were collected as done for the cytokine array, and all of the gels that were probed using antibodies were compared to a control gel that was stained with Sypro Ruby (Invitrogen). The Sypro stained gel served as a loading control for normalization of the western blots and was quantified using ImageJ. Proteins were transferred to a nitrocellulose membrane at 20 V for 2 hours followed by 70 v for an additional hour. The membrane was blocked using 10% milk in TBST rocking at room temperature for 1 hour. The membrane was then incubated with the primary antibody according to manufacturer recommended dilutions at 4°C rocking overnight. Following the primary antibody incubation, the membrane was washed in TBST every 5 minutes for 45 minutes. For the IL-10, osteopontin, and I-309 anti-rabbit secondary antibody at 1:5000 dilution (Santa Cruz Biotechnology sc-2357) was used. For MIP-1δ, anti-goat HRP 1:5000 dilution (Santa Cruz Biotechnology sc-2354) was used. In all cases, the secondary antibody was in 10% milk TBST and was allowed to rock at room temperature for 1 hour. Membranes were then washed for 1 hour with TBST changed every 5 minutes. ECL chemiluminescence reagent (KPL) was then added to membrane according to manufacturer recommendations and imaged on Azure Biosystems c600. Following imaging, the western blots were normalized to the Sypro gel to account for any protein loading differences, and quantified in ImageJ.

### Mouse infections

Mice infections were conducted as in Chua et al 2011 [29]. All experiments were conducted at Indiana University School of Medicine Gary in the laboratory of Dr. Taeok Bae [59]. 5-week-old female Balb/Cj mice were obtained from the Jackson Laboratory and kept 5 or 6 per cage at 23°C with 12 h/ 12 h day/night cycle in the animal facility. Food and water were provided ad libitum throughout the experiment. When the mice had aged to 6 weeks the right flank hair was removed, sprayed with 70% ethanol, and 10^8^ CFU of wt, Δ*sagB*, or PBS in 50 µL sterile PBS were injected into 11 mice/treatment condition. Mice were monitored for 5 days post-infection and lesion size later quantified in ImageJ (NIH). Mice were also weighed daily. During the course of infection, 2 mice from wt and 1 mouse from Δ*sagB* groups were prematurely euthanized by CO2 asphyxiation due to their severe signs of acute disease such as impaired mobility and hunched posture. Following the 5-day time course, all mice were euthanized by CO2 asphyxiation, and the lesion was excised to allow for enumeration of CFUs.

## Declarations

### Ethics Approval and Consent to Participate

The animal experiment was performed by following the Guide for the Care and Use of Laboratory Animals of the National Institutes of Health. The animal protocol was approved by the Institutional Animal Care and Use Committee of Indiana University School of Medicine-Northwest (protocol no. NW-43). Every effort was made to minimize suffering of the animals.

### Consent for Publication

Not applicable

### Availability of Data and Materials

The datasets used and/or analyzed during the current study are available from the corresponding author on reasonable request.

### Author Contributions

TK and SWL designed and conceptualized the study; SWL, FJC, TB, and VAP were involved in supervision of the study; TK, KC, YB, WY, CP, FRF, HMV, DEH, and JNR acquired and analyzed the data. TK and KC wrote the manuscript and SWL assisted in the critical reading and revision of the manuscript. All authors have read and approved the final version of the manuscript.

## Acknowledgments

Thank you to all the members of the Lee lab for comments on the manuscript.

## Additional Files

Additional File 1: Figure S1: (A). Growth curve of wildtype JKD6159 and ΔsagB JKD6159 in TSB broth. Representative curve of biological triplicates. (B). Growth curve of JKD6159 wt and ΔsagB in cell culture media representative for use in cytotoxicity assays. Graph represents one of three biological replicates.

Additional File 2: Table S1: Results from Kinexus antibody array

Additional File 3: Figure S2: Overlay of wt and ΔsagB protein gels. wt=green, ΔsagB=red. Marked areas indicate spots that reproduced between technical and biological duplicates. Image generated using Delta2D software.

Additional File 4: Table S2: Proteins identified from 2D proteomics.

Additional File 5: Table S3: Primer sets used for gene knockouts and in vitro toxin generation.

