## Supplementary figures and images for "Discovery of genes encoding a Streptolysin S-like toxin biosynthetic cluster in a select highly pathogenic methicillin resistant *Staphylococcus aureus* JKD6159 strain"

### Figure S1

Additional File 1: Figure S1.

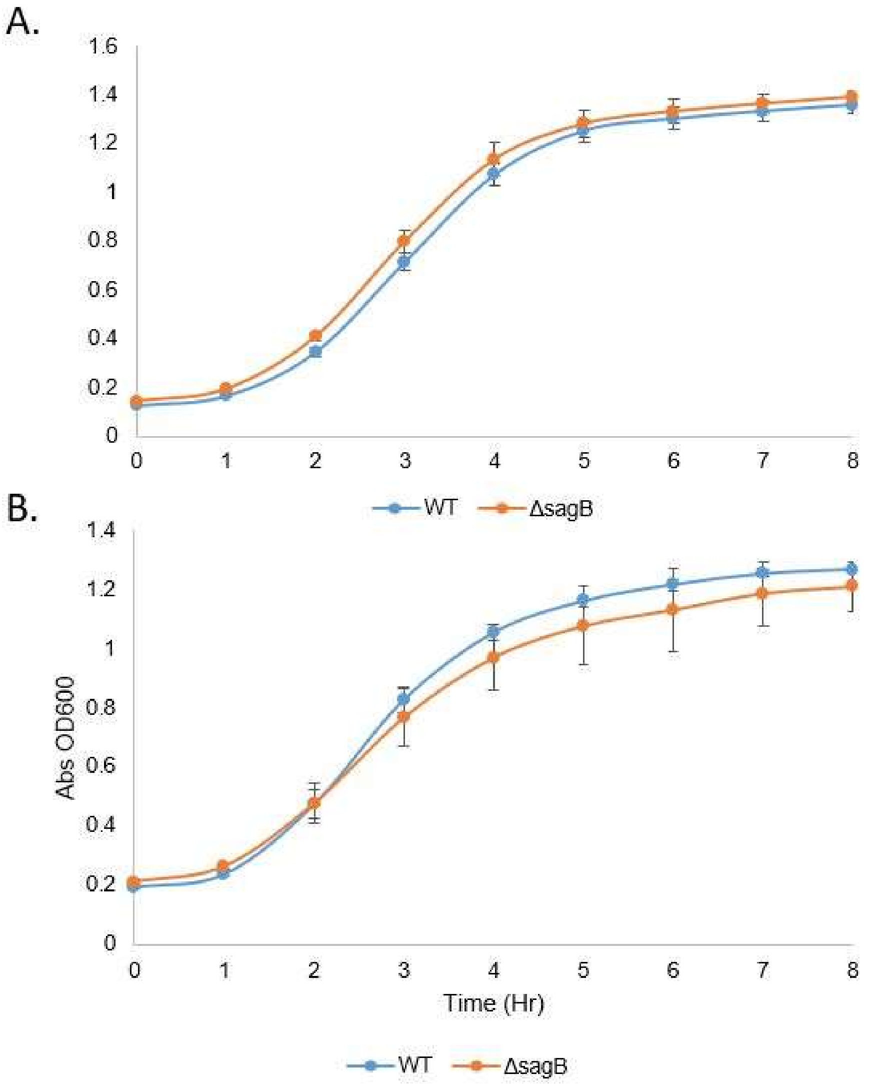

### Figure S2

Additional File 3: Figure S2.

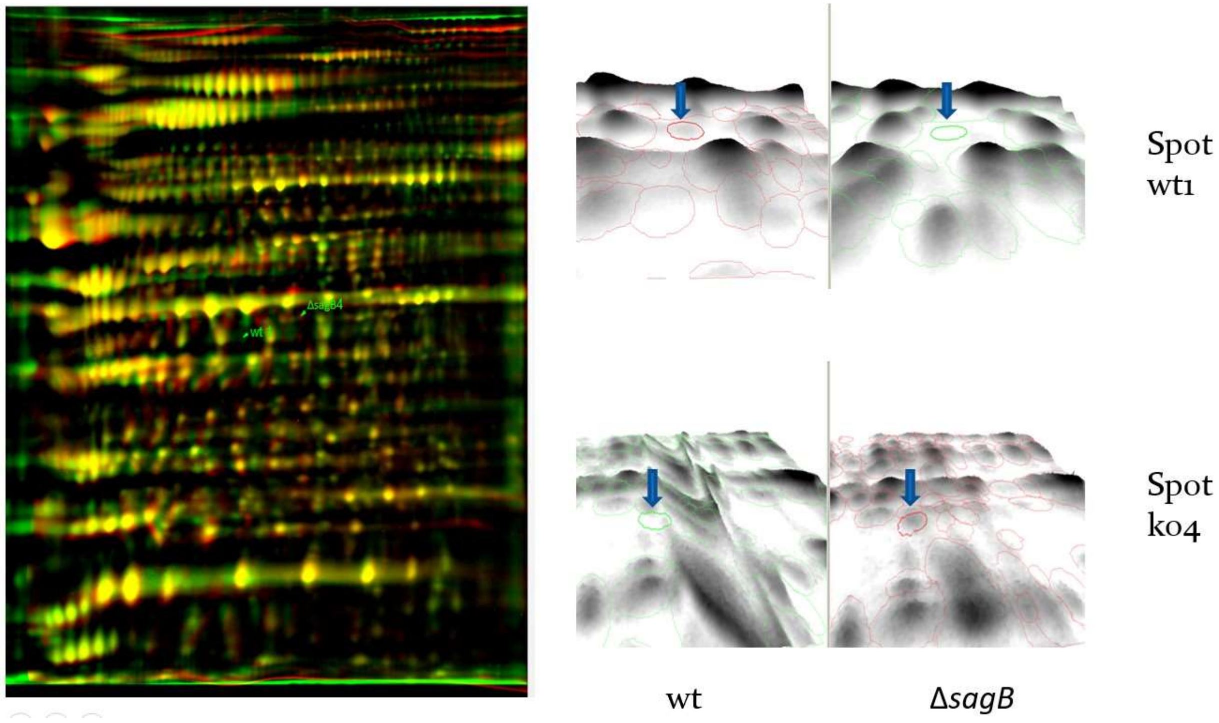
