## Supplementary material for "Discovery of genes encoding a Streptolysin S-like toxin biosynthetic cluster in a select highly pathogenic methicillin resistant *Staphylococcus aureus* JKD6159 strain": Table S1

Additional File 2: Table S1.

| Protein Name | Phosphorylation site | % down regulation in wt |
|---|---|---|
| Cyclin-dependent protein-serine kinase 1 | T14 | 97 |
| Cyclin-dependent protein-serine kinase 1 | T161 | 97 |
| Cyclin-dependent protein-serine kinase 1 | T15/Y15 | 55 |
| Cyclin-dependent protein-serine kinase 1 | Pan-specific | 51 |
| Cyclin-dependent protein-serine kinase 6 | Y13 | 101 |
| Cyclin-dependent protein-serine kinase 6 | Y24 | 65 |
| Cyclin-dependent protein-serine kinase 7 | T170 | 52 |
| Cyclin-dependent protein-serine kinase 9 | T186 | 60 |
| Cyclin-dependent protein-serine kinase 10 | T196 | 64 |
| Epidermal growth factor-tyrosine kinase | Y998 | 85 |
| Epidermal growth factor-tyrosine kinase | T693 | 143 |
