## Supplementary material for "Discovery of genes encoding a Streptolysin S-like toxin biosynthetic cluster in a select highly pathogenic methicillin resistant *Staphylococcus aureus* JKD6159 strain": Table S2

Additional File 4: Table S2.

| **Spot origin** | **Protein Name** | **% coverage** |
|---|---|---|
| Wt 1 | Threonine synthase | 13.60 |
| | 2 oxoisovalerate dehydrogenase subunit alpha | 7.58 |
| | Carbamate kinase | 26.20 |
| | NH3 dependent NAD synthetase | 49.08 |
| | Succinate CoA ligase ADP forming alpha subunit | 45.03 |
| | Thioredoxin disulfide reductase | 36.98 |
| | Putative glucokinase ROK family | 10.98 |
| | D isomer specific 2 hydroxyacid dehydrogenase family protein | 26.90 |
| Δ*sagB* 4 | Uroporphyrinogen decarboxylase | 12.17 |
| | FolD | 38.47 |
| | Global transcriptional regulator catabolite control protein A | 46.50 |
| | Hydroxymethylbilane synthase | 11.36 |
| | FAD dependent pyridine nucleotide disulphide oxidoreductase | 9.45 |
| | S adenosyl methyltransferase MraW | 15.11 |
