## Supplementary material for "Discovery of genes encoding a Streptolysin S-like toxin biosynthetic cluster in a select highly pathogenic methicillin resistant *Staphylococcus aureus* JKD6159 strain": Table S3

Additional File 5: Table S3.

| Primer Name | Primer Sequence | Restriction Enzyme Site (Red) |
|---|---|---|
| sagB KO downF | ATAGGTACCTGAGACGCTGAAGCGATTAAAATATT | Kpn1 |
| sagB KO downR | ATAGAATTCTCATTGCTTTTTACCTCTTTTGTTAGT | EcoR1 |
| sagB KO upF | ATAGAATTCTTGAGCAATGAAAACGTTTATTAAACC | EcoR1 |
| sagB KO upR | ATAGCGGCCGCATGTGGGTAAATAAAAATGAAGTACATTTC | Not1 |
| BF KO chromosome | CCAATTGCTTTTATTAGTTTTCCAATTTG | |
| BR KO chromosome | ACATGTCGAACAAGTAGACTGTC | |
| pIMAYF | GCTTTGGCAGTTTATTCTTGACATGTA | |
| pCL55 insertF | CTTATTTTTAAATTTTTCAAACCACATTTT | |
| pCL55insertR | CTTTCGTCTTCAAGAATTCGAGC | |
| pCL55 sequence F | GACAACACTTACACGTTTCCATTT | |
| pCL55 sequence R | CCACCTGACGTCTAAGAAACC | |
